# The ubiquitin-proteasome system regulates the formation of specialized ribosomes during high salt stress in yeast

**DOI:** 10.1101/2024.08.15.608112

**Authors:** Yoon-Mo Yang, Katrin Karbstein

## Abstract

Rps26-deficient ribosomes are a physiologically relevant ribosome population which arises during osmotic stress to support the translation of mRNAs involved in the response to high salt in yeast. They are formed by binding of the chaperone Tsr2 to fully assembled ribosomes to release Rps26 when intracellular Na^+^ concentrations rise. Tsr2-mediated Rps26 release is reversible, enabling a rapid response that conserves ribosomes. However, because the concentration of Tsr2 relative to ribosomes is low, how the released Rps26•Tsr2 complex is managed to allow for accumulation of Rps26-deficient ribosomes to nearly 50% of all ribosomes remains unclear. Here we show that released Rps26 is degraded via the Pro/N-degron pathway, enabling the accumulation of Rps26-deficient ribosomes. Substitution of the N-terminal proline of Rps26 to serine increases the stability of free Rps26, limits the accumulation of Rps26-deficient ribosomes and renders yeast sensitive to high salt. The GID-complex, an E3 ubiquitin ligase, and its adaptor Gid4, mediate polyubiquitination of Rps26 at Lys66 and Lys70. Moreover, this ubiquitination event is required for Rps26 degradation, the accumulation of Rps26-deficient ribosomes and the high salt stress resistance. Together, the data show that targeted degradation of released Rps26 from the Rps26•Tsr2 complex allows Tsr2 to be recycled, thus facilitating multiple rounds of Rps26 release.

## Introduction

Ribosomes function to synthesize proteins within cells. Composed of rRNA and ribosomal proteins, more than half of all transcription and translation processes are devoted to the production of ribosome components and to maintain an adequate number of ribosomes (Shore & Albert 2022, Warner 1999). Correct assembly and maintenance of an appropriate number of ribosomes are both crucial for intracellular protein homeostasis (Mills & Green 2017, Parker & Karbstein 2023). Recent studies, however, have shown that ribosomes can have diverse compositions (Emmott et al 2019, Ferretti & Karbstein 2019, Genuth & Barna 2018a, Genuth & Barna 2018b, Martinez-Seidel et al 2020, Norris et al 2021, Sauert et al 2015). While many of these ribosomes result from haploinsufficiency of ribosomal proteins and are associated with diseases (Blomqvist et al 2023, Bolze et al 2013, Collins et al 2018, Fortier et al 2015, Kondrashov et al 2011), several cases have been found in wild-type cells as well (Ferretti et al 2017, Loveland et al 2016, Samir et al 2018, Shi et al 2017, Slavov et al 2015, Sun et al 2021). One of these altered ribosome populations with known physiological roles are Rps26-deficient ribosomes, which form when yeast cells are exposed to high Na^+^ or high pH stress (Ferretti et al 2017). Rps26 binds the mRNA at the −4 residue (upstream of the start codon) of the Kozak sequence, conferring preference for adenosine. Correspondingly, Rps26-deficient ribosomes enable the translation of mRNAs lacking the preferred adenosine residue at position −4 of the Kozak sequence. This altered mRNA preference allows yeast to translate a subset of mRNAs from the Hog1 or Rim101 pathways, enabling the response to high Na^+^ or high pH stress (Ferretti et al 2017). Formation of Rps26-deficient ribosomes during these stress conditions is managed by the chaperone Tsr2 (Yang & Karbstein 2022), a protein previously suggested to deliver Rps26 to ribosomes (Schutz et al 2014, Schutz et al 2018). Tsr2 releases Rps26 from pre-existing mature ribosomes (**Figure 1A**, (Yang & Karbstein 2022)). Moreover, Tsr2 also promotes the dissociation of Rps26 during oxidative stress by specifically removing oxidatively damaged Rps26 from mature ribosomes, enabling their repair (Yang et al 2023). In both of these cases, Tsr2 also reforms Rps26-containing ribosomes by promoting the reincorporation of Rps26. Binding and release of Rps26 is regulated by its affinity to ribosomes, which is modified by high salt and oxidative damage to the protein. Thus, the formation of Rps26-deficient ribosomes should be sensitive to the amount of the Tsr2•Rps26 complex and ultimately Rps26. Binding to its chaperone Tsr2 renders Rps26 stable even without binding to ribosomes (Schutz et al 2014). Thus, it is likely that Rps26 released under these conditions would be stable with Tsr2. Considering that Tsr2 is essential for the release of Rps26 from ribosomes by forming the Rps26•Tsr2 complex (Yang et al 2023, Yang & Karbstein 2022), and considering that binding and release follow an equilibrium governed by mass-action, a stable Tsr2•Rps26 complex should limit the amount of Rps26-deficient ribosomes to the amount of Tsr2 in the cell. Nonetheless, the intracellular levels of Tsr2 are relatively low compared to ribosomes (~5%), whereas Rps26-deficient ribosomes accumulate up to 50% of total ribosomes (**Figure 1B**, (Yang & Karbstein 2022)), suggesting that Tsr2 must catalyze successive rounds of Rps26 release, which would require release and degradation of Rps26.

**Figure 1:**
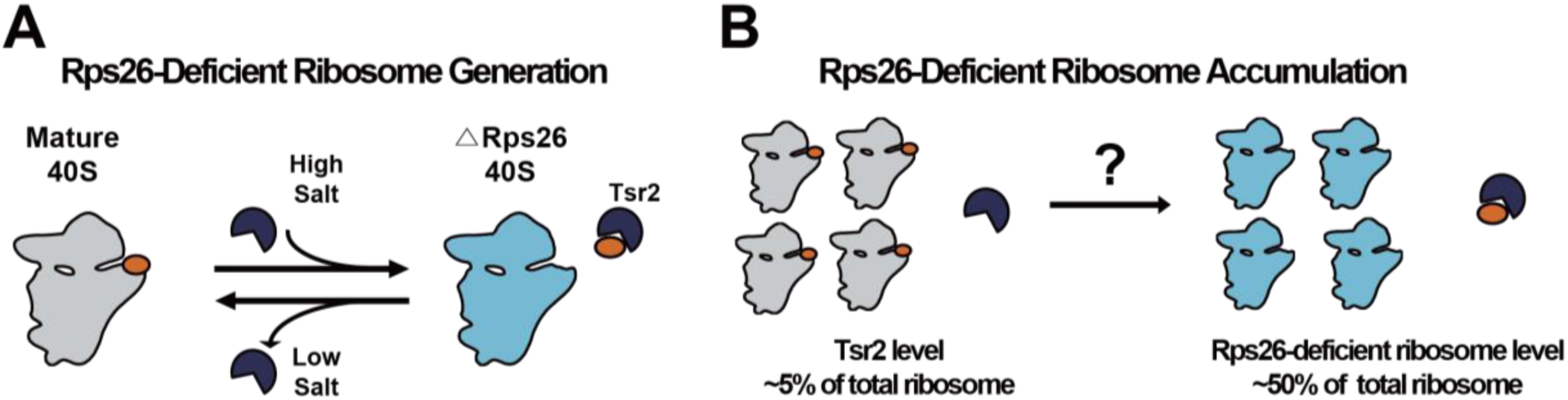
Accumulation of Rps26-deficient ribosome during high salt stress. (A) During high salt stress, the chaperone Tsr2 (dark blue) binds 40S ribosomes (grey) to release Rps26 (orange) and thereby produce Rps26-deficient ribosomes (light blue; (Yang & Karbstein 2022)). This process is reversible when the stress subsides. (B) Rps26-deficient ribosome accumulate up to ~50% of total ribosome (Ferretti et al 2017, Yang & Karbstein 2022). However, levels of Tsr2 are ~5% compared to total ribosome level (Cherry et al 2012).

Because oxidatively damaged Rps26 is degraded after release (Yang et al 2023), we considered whether Rps26 released during high salt or high pH stress is also degraded. This would free Tsr2 for multiple rounds of Rps26 release and remove the released Rps26 from the equilibrium.

Here, we report that the N-terminal proline of Rps26 induces the degradation of released Rps26, which is ultimately important for the re-generation of free Tsr2 to accumulate Rps26-deficient ribosomes. Our results show that replacement of the Rps26 N-terminal proline with serine affects the yield of Rps26-deficient ribosomes under salt stress. Additionally, our data show that the GID-complex and its adaptor subunit Gid4 (Chen et al 2017), a multi-subunit E3 ligase, recognize the N-terminal proline of Rps26 and ubiquitinate Lys66 and Lys70 of Rps26, which eventually leads to degradation of Rps26 under high salt stress. Importantly, this process is required for high salt resistance in yeast.

## Results

### Rps26_P2S limits the accumulation of Rps26-deficient ribosomes under high salt stress

In considering whether Rps26 outside of the ribosome could be selectively degraded to accumulate Rps26-deficient ribosomes, we considered the well-known degradation of proteins via the so-called N-degron pathway. Certain N-terminal residues mediate selective ubiquitination, which render the protein unstable (Varshavsky 2019). One such destabilizing amino acid is proline (Chen et al 2017), which is the N-terminal amino acid in Rps26 (after co-translational removal of the methionine start codon). Moreover, the N-terminus of Rps26 is buried inside the ribosome (**Figure 2A-B**), providing for a possible mechanism for selective destabilization of Rps26 outside the ribosome. We therefore hypothesized that release of Rps26 from the ribosome would uncover the N-terminal proline and lead to degradation of the protein. To test this hypothesis, we utilized the previously identified Rps26 mutant Rps26^D33N^. This mutation weakens its binding to the ribosome and leads to the accumulation of ribosomes lacking Rps26 (Schutz et al 2014, Yang & Karbstein 2022). To trace the degradation of pre-existing Rps26, we designed a pulse-chase experiment where pre-existing Rps26 was marked with an HA-tag whose plasmid-based expression relies on a galactose-inducible/glucose-repressible promoter. In addition, Rps3-HA is also expressed from a galactose-inducible/glucose-repressible promoter. Cells were initially grown in galactose media, such that a portion of pre-made Rps26 and Rps3 will be HA-tagged. In mid log phase, cells were switched to glucose media, where only endogenous untagged ribosomal proteins are expressed, allowing us to trace the pre-existing HA-tagged Rps26 and Rps3 using anti-HA Western blotting. Control experiments confirm that pre-existing Rps26-HA and Rps3-HA are stable over time when they are bound to ribosomes (**Figure 2C-D**). However, with the weak binding Rps26^D33N^, we observed a rapid degradation of Rps26 with a half-life ~30 min, while Rps3-HA remains stable (**Figure 2C-D**). Since free ribosomal proteins are already known to degrade rapidly via the E3 ligase Tom1 (Gorenstein & Warner 1977, Sung et al 2016, Warner 1977), we further tested whether the degradation rate of free-Rps26 is additionally affected by the N-terminal proline. For this we generated a weak binding Rps26 mutant presenting N-terminal serine instead of proline (Rps26^P2SD33N^). Importantly, free-Rps26 harboring an N-terminal serine was more stable than Rps26 bearing an N-terminal proline with increased half-life of ~90 min (**Figure 2C-D**). As described above, in yeast that experience high salt stress, Tsr2 releases Rps26 from ribosomes, producing Rps26-deficient ribosomes that make up nearly half of all ribosomes. Given that Tsr2 levels are ~5% of ribosome levels, we hypothesized that degradation of Rps26 was required for repeated cycles of Rps26 release and the accumulation of Rps26-deficient ribosomes to nearly 50%.

**Figure 2:**
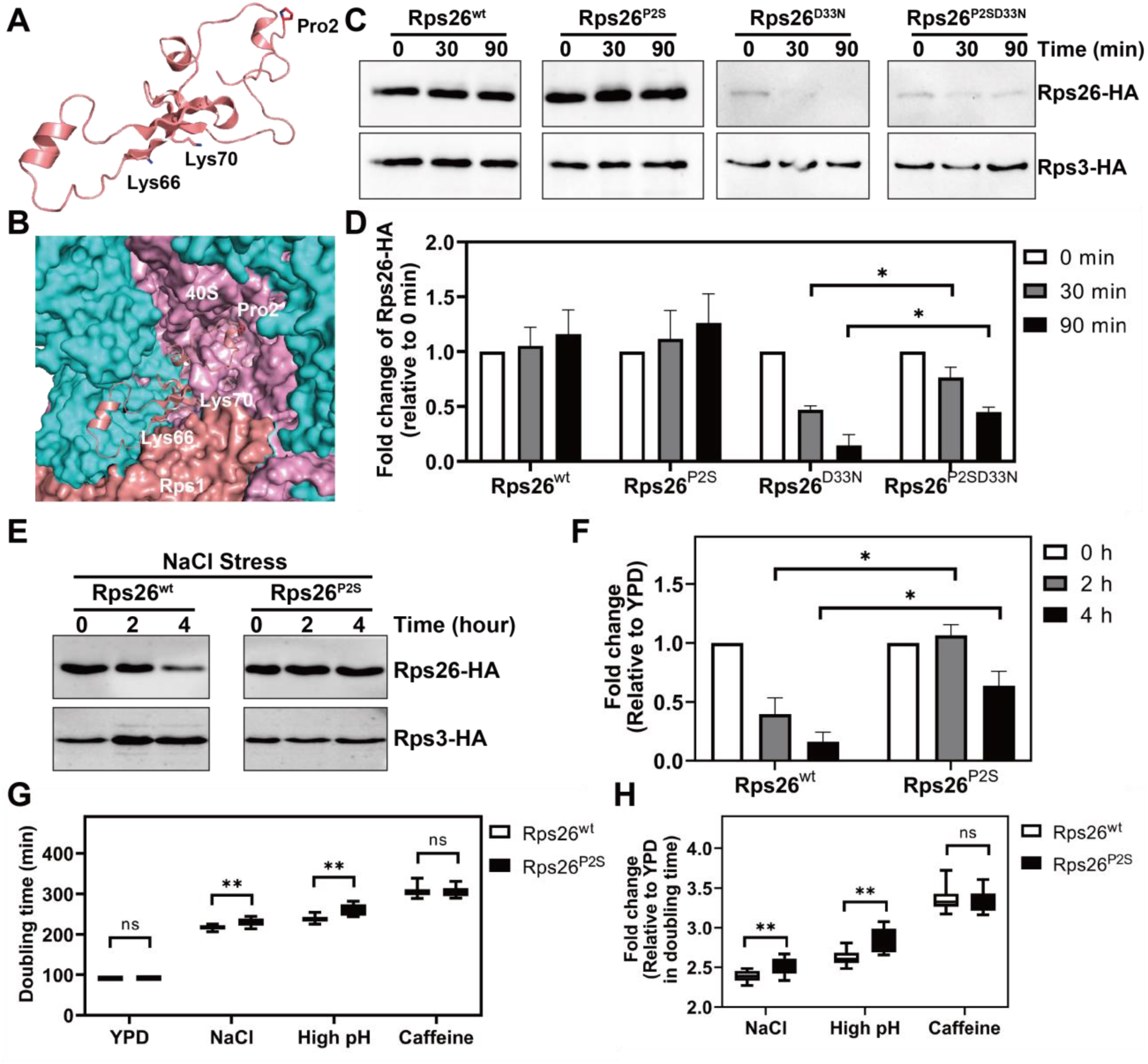
The N-terminal proline of Rps26 is important for accumulation of Rps26-deficient ribosomes during high salt stress. (A-B) The position of proline 2 in Rps26 alone (A) or bound to the 40S subunit (B, PDB 4V88). 18S rRNA is colored in pink, Rps1 colored in salmon and other ribosomal proteins colored in cyan. (C) Western blot to measure degradation of pre-existing Rps26 and Rps3 over time. Pulse-chase experiments to separate pre-existing Rps26-HA and Rps3-HA from newly-made ribosomal proteins Rps26 and Rps3 rely on a wild type yeast strain (BY4741) with Rps26-HA (pkk30528) and Rps3-HA (pKK31042) produced from plasmid-based galactose-inducible/glucose-repressible promoters. By shifting this strain from galactose to glucose, pre-existing ribosomal proteins are marked with the HA-tag for further detection. (D) Quantification of Rps26-HA stability relative to Rps3-HA stability data in panel (C). Data are averaged from 3 biological replicates. Error bars represent the SEM, and significance was determined using an unpaired t-test. *p < 0.05. (E) Western blot to measure degradation of pre-existing Rps26-HA and Rps3-HA during NaCl stress. By shifting wt cells containing pkk30528 (or pKK31150) and pKK31042 from galactose to glucose with 1 M NaCl, degradation of pre-existing ribosomal proteins was traced via their HA-tag. (F) Quantification of data in panel (E). Data are averaged from 3 biological replicates. Error bars represent the SEM, and significance was determined using an unpaired t-test. *p < 0.05. (G) Changes in doubling time upon addition 1 M NaCl or high pH (8.2) or caffeine, in YKK491 cells (Gal:Rps26) with Rps26 variants produced from plasmids driven by the TEF promoter. Data are the average of 3 biological replicates and 3 technical replicates. **p < 0.01 by unpaired t-test. (H) Values in (G) normalized to non-stress conditions (fold change = 1). **p < 0.01 by unpaired t-test.

We thus wanted to next test if the N-proline-driven degradation of Rps26 was required for the accumulation of Rps26-deficient ribosomes under conditions of high salt stress (Ferretti et al 2017). We therefore measured the stability of pre-existing HA-tagged Rps26^wt^ and Rps3 after high salt treatment, which leads to Tsr2-mediated release of Rps26 from ribosomes. The data in **Figure 2E-F** demonstrate that increased concentrations of NaCl reduce the stability of pre-existing wt Rps26-HA relative to Rps3-HA, as expected from the reduced occupancy of Rps26 in ribosomes under these conditions (Ferretti et al 2017, Yang & Karbstein 2022). However, substitution of the N-proline to serine stabilized Rps26 after NaCl treatment (**Figure 2E-F**). Thus, the N-proline was required for the degradation of Rps26 during high salt stress.

Rps26-deficient ribosomes are required for resistance to high salt and high pH, but not caffeine (Ferretti et al 2017, Yang & Karbstein 2022). Thus, to further test whether mutation of the N-proline blocked the accumulation of ribosomes lacking Rps26, we measured the sensitivity of yeast expressing Rps26^wt^ or Rps26^P2S^ to high salt, high pH and caffeine. As predicted, relative to Rps26^wt^, Rps26^P2S^ was sensitive to high salt and high pH, but not caffeine (**Figure 2G-H**). These data strongly suggest that indeed degradation of Rps26 via the N-proline is required for the accumulation of Rps26-deficient ribosomes and demonstrate a physiological role for this process.

### Rps26 Lys66 and Lys70 are ubiquitinated under high salt stress and are important for degradation of Rps26

We next wanted to know if ubiquitination was part of the mechanism for decay of released Rps26 as for other proteins of the N-end pathway (Tasaki et al 2012). Therefore, we individually mutated the five known (Swaney et al 2013) ubiquitinated lysine residues in Rps26 to arginine and tested whether any mutation affects the degradation of Rps26 under high salt stress (**Figure 3A-B**). By tracing the pre-existing Rps26-HA variants, we observed increased stability of Rps26 when either Lys66 or Lys70 were mutated to arginine. Other mutants (K28R, K108R and K116R) showed similar degradation as Rps26^wt^ suggesting their irrelevance for Rps26 degradation. Interestingly, Lys66 and Lys70 residues are protected by Rps1 and 18S rRNA, respectively, but become surface exposed after release from ribosome (**Figure 2A-B**). Thus, ubiquitination on these sites would only occur when Rps26 dissociates from ribosomes. Moreover, both residues are located in the domain that interacts with Tsr2 (Schutz et al 2018), indicating that ubiquitination of these sites could weaken the interaction with Tsr2 (See discussion).

**Figure 3:**
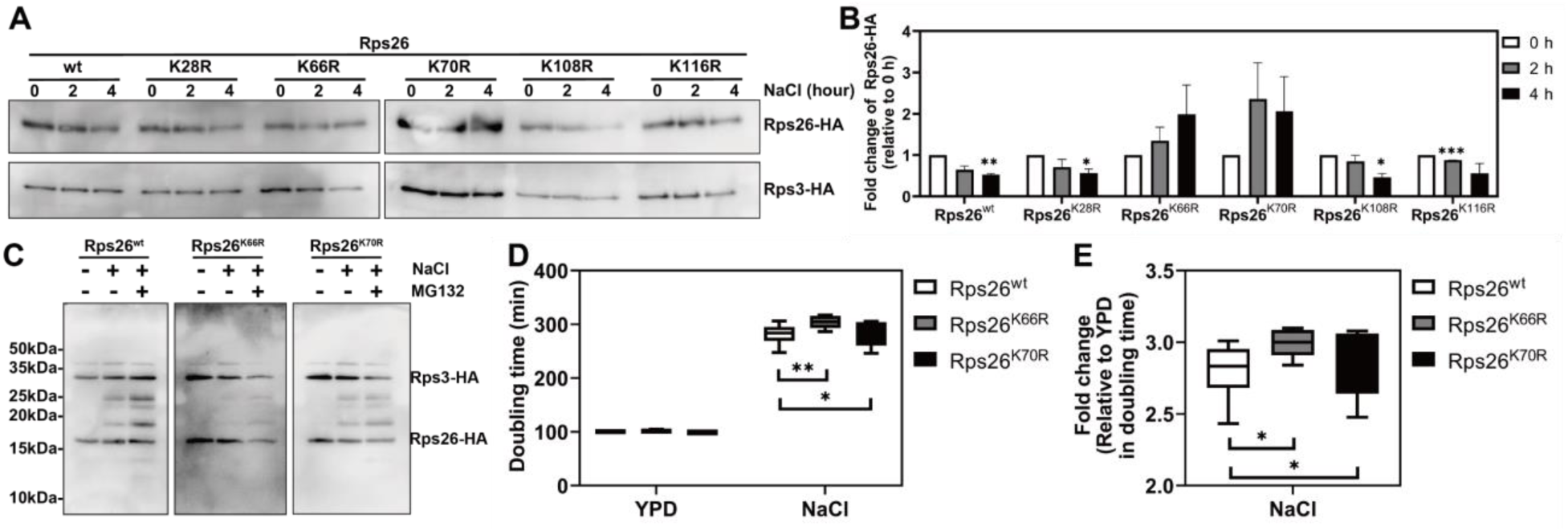
Lys66 and Lys70 of Rps26 are ubiquitinated during high salt stress. (A) Western blot to measure degradation of pre-existing Rps26-HA variants and Rps3-HA. By shifting cells from galactose to glucose with 1 M NaCl, degradation of pre-existing Rps26 variants were tranced with the HA-tag. (B) Quantification of data in panel (A). Data are averaged from 2 biological replicates. Error bars represent the SEM, and significance was determined using an unpaired t-test. *p < 0.05, **p < 0.01, ***p < 0.001, by unpaired t-test. (C) Western blot to measure ubiquitination of pre-existing Rps26-HA variants. By shifting cells from galactose to glucose with 1 M NaCl, cells were treated or untreated with 5 µM of MG132 for 12 h before collection. (D) Changes in doubling time upon addition 1 M NaCl, in YKK491 cells (Gal:Rps26) with Rps26 variants produced from plasmids driven by the TEF promoter. Data are the average of 3 biological replicates and two technical replicates. *p < 0.05, **p < 0.01 by unpaired t-test. (E) Values in (D) normalized to non-stress conditions (fold change = 1). *p < 0.05 by unpaired t-test. (F) Model for Rps26-deficient ribosome accumulation by ubiquitin-proteasome system under high salt stress.

To further confirm directed ubiquitination of these two residues, we collected cells after treatment with NaCl and the proteasome inhibitor MG132, to block the degradation of ubiquitinated Rps26, and thus enable its accumulation. As expected, higher molecular weight bands of wt Rps26 increase with high salt treatment and these variants increase even more after treatment with MG132 (**Figure 3C**). Importantly, when Lys66 was mutated to arginine, accumulation of ubiquitinated Rps26 significantly decreased, further supporting Lys66 as a site of ubiquitination during high salt stress. Mutation of Lys70 also reduced the amount of ubiquitinated Rps26, but somewhat less than K66R, indicating that it can also be targeted by the GID complex, as expected based on stabilization of Rps26 by its mutation to arginine.

To further confirm K66 and K70 as sites of ubiquitination and verify that ubiquitination (and subsequent decay) of Rps26 is important for high salt stress resistance, we measured the growth effect from high salt treatment in cells expressing Rps26^wt^, Rps26^K66R^ or Rps26^K70R^ (**Figure 3D-E**). While mutation of Lys66 or Lys70 had no effect under normal growth conditions, both K66R and K70R showed increased sensitivity to high salt stress compared to Rps26^wt^. Together these results suggest that ubiquitination of Rps26 on both K66 and K70 are important for its decay, the generation of Rps26-deficient ribosomes, and thus the resistance to high salt stress.

### Degradation of Rps26 is affected by recognin Gid4 of the GID-complex

Previous work has shown that some N-terminally destabilizing amino acids are recognized by the GID complex, a multi-protein E3 ubiquitin-ligase, which uses specific adapters, referred to as recognins, to identify specific N-terminally destabilizing sequences (Chen et al 2017). Moreover, Gid4 and Gid10 have been identified as N-proline recognizing adaptors that are induced under osmotic stress (Chen et al 2017, Langlois et al 2022, Melnykov et al 2019). We therefore assessed the stability of released Rps26 in yeast strains lacking Gid4 or Gid10. Importantly, the data in **Figure 4A-B** show that in cells lacking Gid4, but not in cells lacking Gid10, Rps26-HA is stabilized under high salt stress relative to cells containing Gid4. Control experiments confirm that Rps26 is stable in this strain when the N-proline is substituted with serine (**Figure 4C-D**). These data demonstrate that Gid4 is important for the degradation of Rps26 via the Pro/N-degron dependent pathway.

**Figure 4:**
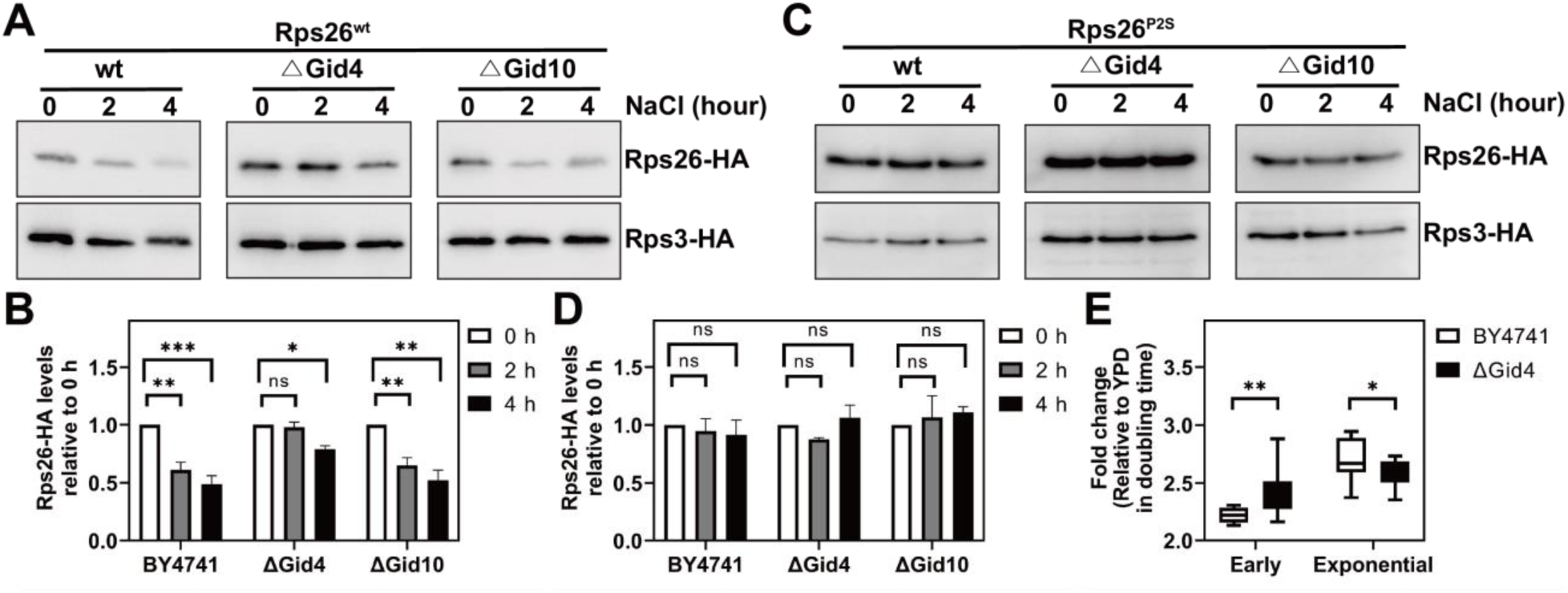
Gid4 is important for the accumulation of Rps26-deficient ribosomes. (A, C) Western blot to measure degradation of pre-existing Rps26-HA and Rps3-HA in the presence and absence of Gid4 and Gid10. By shifting wt or ΔGid4 cells expressing wt Rps26^wt^-HA (A), or Rps26^P2S^-HA (C) and Rps3-HA from galactose to glucose with 1 M NaCl, degradation of pre-existing ribosomal proteins was traced via the HA-tag. (B) Quantification of data in panel (A). Data are averaged from 2-4 biological replicates. Error bars represent the SEM, and significance was determined using an unpaired t-test. *p < 0.05. (D) Quantification of data in panel (C). Data are averaged from 2 biological replicates. Error bars represent the SEM. (E) Changes in doubling time upon addition 1 M NaCl in wt or cells lacking Gid4. Data are the average of 4 biological replicates and 3 technical replicates and were normalized to non-stress conditions (fold change = 1). *p < 0.05, **p < 0.01 by unpaired t-test.

To further confirm that Gid4-mediated decay of Rps26 is required for the accumulation of Rps26-deficient ribosomes, which mediate the resistance of yeast to high salt (Ferretti et al 2017), we again measured the sensitivity of yeast to grown in high salt media. Indeed, Gid4 is important for the recovery from high salt stress (**Figure 4E**). Surprisingly, Gid4 is not required after ~ 400 minutes, when cells re-enter exponential growth (**Figure 4E**). This contrasts with the requirement for the N-proline, which persists into exponential phase (**Figure 2G-H**) and indicates that a second recognin is required later in exponential phase. This may also explain why the effect caused by proline substitution is greater than deletion of the recognin since no alternative recognition would occur by N-terminal serine (see discussion).

## Discussion

### Released Rps26 is degraded via the N-degron pathway, enabling multiple cycles of Rps26 release

Yeast exposed to high salt (or pH) stress release Rps26 from about half of their ribosomes, in a process that is mediated by the chaperone Tsr2 (Ferretti et al 2017, Yang & Karbstein 2022). Because Tsr2 levels are about 5% of ribosome levels, but Rps26-deficient ribosomes make up nearly 50% of all ribosomes when yeast experience high salt stress, each Tsr2 must release ~ 10 Rps26 from ribosomes. How the released Rps26 is removed from the Tsr2•Rps26 complex (without allowing it to rebind) to enable a new round of Rps26 release was unknown. Here, we how Tsr2 can mediate repeated cycles of Rps26 release due to the specific degradation of the released Rps26 via the ubiquitin proteasome system (**Figure 5A**).

**Figure 5:**
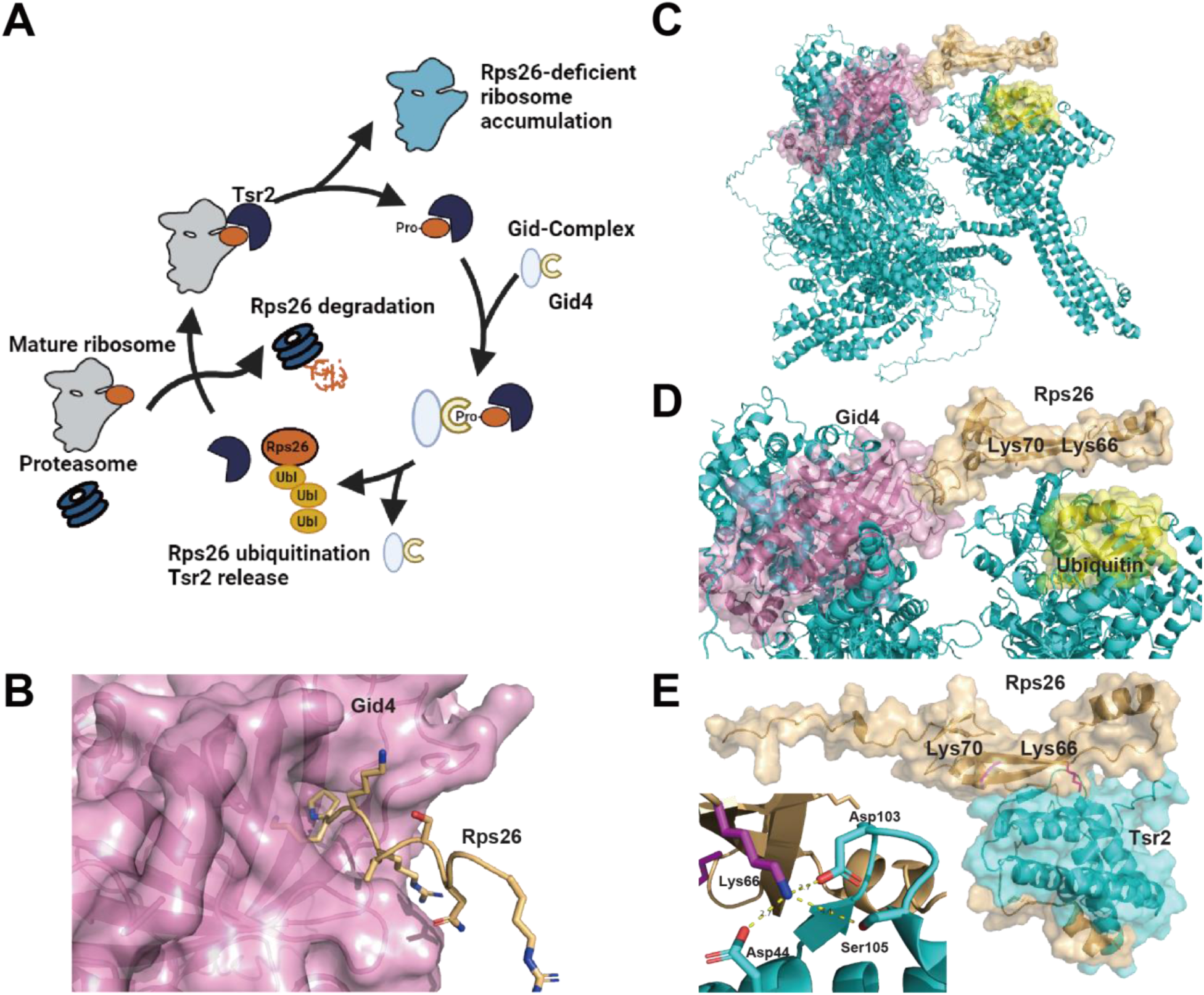
Gid4-mediated ubiquitination and decay of Rps26 enables repeated cycles of Rps26 release by Tsr2. (A) Model for accumulation of Rps26-deficient ribosomes via repeated cycles of Tsr2-mediated Rps26 release enabled by the ubiquitin-proteasome system during high salt stress. During high salt stress, the chaperone Tsr2 (dark blue) binds mature 40S ribosomes (grey) to release Rps26 (orange) and thereby produce Rps26-deficient ribosomes (light blue). The N-terminal proline on the released Rps26 is recognized by the GID complex (grey) and the Gid4 adaptor (yellow), enabling ubiquitination and ultimately proteasome-mediated decay of the released Rps26, and freeing Tsr2 for another round of Rps26 release. (B) AlphaFold (Jumper et al 2021) structural prediction of Gid4 (pink) bound to the Rps26 N-terminal peptide (P^2^KKRASNGER^10,^ shown in orange). (C-D) AlphaFold-generated structural prediction of Rps26 bound to the GID-complex. AlphaFold was used to predict the structure of the Gid-complex (Gid1, Gid2, Gid3, Gid5, Gid8, Gid9, cyan), bound to Gid4 (pink) and ubiquitin (yellow) with the N-terminal peptide (P^2^KKRASNGER^10^) of Rps26. Then Rps26 (orange, using amino acid 2-87) was aligned to the predicted GID-substrate-complex using the N-terminal peptide as a reference. (E) AlphaFold-generated structural prediction of the Tsr2-Rps26 complex. Rps26 and Tsr2 are represented in orange and cyan, respectively. Lys66 of Rps26 interacts with Asp44, Asp103 and Ser105 of Tsr2, consistent with biochemical data (Schutz et al 2018).

Our data show that the N-terminal proline of Rps26 affects the stability of Rps26 when released from the 40S subunit. Importantly, N-proline-mediated decay of Rps26 is required for the accumulation of Rps26-deficient ribosomes and the response to high salt stress. After Rps26 is released, Gid4, the adaptor protein of the GID E3 ubiquitin ligase complex gets access to the N-terminal proline of Rps26. This allows the GID-complex to recognize Rps26 as a target protein and subsequently ubiquitinate Rps26 for selective degradation. Importantly, this pathway is distinct from the much slower degradation of unincorporated ribosomal proteins that serves to maintain the stoichiometry of ribosomal proteins and is mediated by Tom1.

While in most cases it is unclear how these Pro/N-end substrates are specifically degraded under certain conditions, our results imply that for Rps26 temporal regulation arises via the accessibility of the N-terminus of Rps26 (**Figure 2A-B**). Rps26 is stable when the N-terminal proline is buried inside the ribosome complex, which prevents Rps26 from being recognized by Gid4. Indeed, AlphaFold (Jumper et al 2021) predicts an interaction between the N-terminus of Rps26 and Gid4 (**Figure 5B**), consistent with these biochemical data.

### Ubiquitination occurs in the Tsr2 recognition site

Ubiquitination of Lys66 and Lys70 is critical for accumulating Rps26-deficient ribosomes (**Figure 3**). Excitingly, when we use AlphaFold (Jumper et al 2021) to predict the structure of the GID-complex bound to ubiquitin and the N-terminus of Rps26, and then align full-length Rps26 with the N-terminus by overlaying the identical amino acids, the resulting GID-Rps26 complex places Lys70, and also Lys66 adjacent to the ubiquitin, consistent with the biochemical data (**Figure 5C-D**).

Structural prediction using AlphaFold (Jumper et al 2021) suggests that Lys66 and Lys70 of Rps26 are positioned at the interface with Tsr2 (**Figure 5E**), consistent with previous biochemical data (Schutz et al 2018). Specifically, Lys66 is predicted to form a charge-charge interaction with Tsr2. Ubiquitination at this site would likely disrupt this charge interaction, in addition to causing structural hindrance between the two proteins. Thus, ubiquitination is expected to release Rps26 from Tsr2, leading to the degradation of Rps26 alone, and enabling Tsr2 to carry out another cycle of Rps26 release.

### Do other E3 ligases impact Rps26 stability?

Interestingly, our results show that Gid4 is important for the degradation of Rps26 during the early phase of the stress response, while Gid4 is not required for prolonged growth in high salt (**Figure 4E**). This suggests that other E3 ligases might complement the role of Gid4 later in the adaptation. Indeed, we observed a slight decrease in Rps26 levels in Gid4 deletion strains after 4 hours of stress treatment (**Figure 4A-B**). Moreover, unlike Gid4, which shows basal expression under normal conditions, Gid10 expression is tightly repressed in YPD and induced after salt stress (data not shown, (Langlois et al 2022)), suggesting an alternative degradation pathway of Rps26 caused by Gid10, the other known proline-recognizing E3 ligase (Langlois et al 2022, Melnykov et al 2019). While we were not able to detect an impact from Gid10 on Rps26 stability (**Figure 4**), it is possible that complex feedback regulation and/or small effect sizes might have masked such an affect, which would require additional experimentation.

### Are there other examples of chaperone/proteasome-dependent ribosome remodeling?

Ribosome specialization, whereby ribosomes of different composition preferentially translate different subsets of mRNAs is an exciting emerging concept (Emmott et al 2019, Gay et al 2022, Joo et al 2022, Li & Wang 2020, Martinez-Seidel et al 2020, Mauro & Edelman 2002, Norris et al 2021, Sloan et al 2017, Xue & Barna 2012). While such specialization could potentially represent another ubiquitous layer in regulated gene expression, the concept has also significant conceptual flaws (Barna et al 2022, Ferretti & Karbstein 2019, Mills & Green 2017). One of these is that ribosomes are exceptionally long-lived, much longer than the doubling times for most cell types (Yang & Karbstein 2024). Moreover, stress conditions, which often necessitate a change in the gene expression program, also generally downregulate ribosome assembly (Gasch et al 2000, Warner 1999). Thus, degrading existing ribosomes and making new ribosomes would take too long to be relevant on a biological timescale. Remodeling of pre-existing ribosomes presents a potential solution for this conceptual problem. Indeed, we have recently shown that chaperones specialized for individual ribosomal proteins can release Rps26 and Rpl10 from their subunits, either when they are oxidatively damaged (Yang et al 2023), or during high salt or pH stress (Yang & Karbstein 2022). At least 13 ribosomal proteins have personalized chaperone proteins (Pillet et al 2017, Yang & Karbstein 2024). Moreover, with one exception the targets of these chaperones are all located on the outside of the ribosome (Yang & Karbstein 2024), raising the potential that chaperone-mediated ribosome-remodeling could be more common. Nonetheless, none of these candidates have traditionally characterized N-degron signals.

Five additional ribosomal proteins contain an N-terminal proline in yeast, including Rps9, Rps19, Rps25, Rpl12 and Rpl28 (Chen et al 2017). While none of these none have a described chaperone, AROS has been suggested as a chaperone for human Rps19 (Singh et al 2021). Moreover, Rps19 is exchangeable (Yang & Karbstein 2024). Interestingly, yeast express two Rps19 paralogs. While Rps19A contains the N-terminal proline, this residue is substituted to alanine in Rps19B. Despite the highly conserved sequence identity of the yeast ribosomal protein paralogs, some studies have shown the importance of paralog substitutions during stress conditions (Ghulam et al 2020, Malik Ghulam et al 2022), which in some cases can be traced back to differential changes in the expression of the two paralogs under different conditions.

Together, this suggests the possibility that chaperone-mediated release and subsequent degradation of Rps19A but not Rps19B might be a mechanism to produce Rps19-deficient ribosomes in the precise stoichiometry dictated by the relative abundance of Rps19A and Rps19B.

## Materials and Methods

### Strains and plasmids

*S. cerevisiae* strains used in this study were either purchased from the GE Dharmacon Yeast Knockout Collection or constructed using standard methods (Longtine et al 1998) and are listed in Table S1. Plasmids are listed in Table S2.

### Degradation of Rps26

Degradation of Rps26 was tested by growing BY4741 cells transformed with pKK30528 (Gal::Rps26-HA) and pKK31042 (Gal::Rps3-HA) at 30°C to mid-log phase in galactose minimal media. Cells were washed twice and then transferred to YPD containing 0 or 1M NaCl for the indicated time, collected and analyzed by western blotting. To detect ubiquitinated Rps26, cells were initially cultured in in galactose minimal media and transferred to YPD containing 1M NaCl with or without addition of 5 µM MG132 (Sigma). After 12 hours, cells were harvested, washed and then resuspended in 500 µl of lysis buffer (20 mM Hepes/KOH (pH 7.4), 100 mM KOAc, 2.5 mM Mg(OAc)_2_) supplemented with 1 mg/ml heparin, 1 mM benzamidine, 1 mM PMSF and complete protease inhibitor cocktail (Roche). After lysis, each sample was layered over 50 μl of sucrose cushion ((20 mM Hepes/KOH (pH 7.4), 100 mM KOAc, 500 mM KCl, 2.5 mM Mg(OAc)_2_, 1 M Sucrose, 2 mM DTT) and spun in a Beckman TLA 100.1 rotor for 70 min at 400,000 × g. The pellet was resuspended in storage buffer (lysis buffer, 250 mM sucrose and 2 mM DTT) and analyzed by SDS-PAGE followed by Western blotting.

### Quantitative yeast growth measurements

Gal::Rps26 cells (YKK491) supplemented with plasmids encoding the Rps26 variants were grown in appropriate glucose minimal media overnight, and then diluted into fresh YPD for ~2 hours before inoculating into 96-well plates (Thermo Scientific) at a starting OD_600_ between 0.04 and 0.1. A Synergy.2 plate reader (BioTek) was used to record the OD_600_ for 24 hours, while shaking at 30°C. The composition of stress-media was YPD + 1 M NaCl, or YPD + 100 mM TAPS (pH 8.2), or YPD + 10 mM caffeine. Doubling times were calculated using data points within the early phase or mid-log phase using GraphPad Prism 9. Statistical analyses for each measurement are detailed in the respective figure legend.

### Western analyses and Antibodies

Western blots were scanned using the ChemiDoc MP Imaging System from Biorad after applying luminescence substrates (Invitrogen) and quantified using its built-in image lab software (ver. 6.0.1). Intensity of each band was analyzed after local background subtraction. To detect HA-tagged proteins anti-HA antibody from Abcam (ab18181) was used.

## Acknowledgements

We thank members of the Karbstein laboratory for discussion and comments on the manuscript. This work was supported by NIH grant R35-GM136323 to K.K., and by the National Research Foundation of Korea (NRF) grant funded by the Korea government (MSIT) RS-2024-00349721, and the research fund of Hanyang University HY-202300000003039 to Y.Y.

---

**Table S1:** Yeast strains used in this work.

| Strain | Description | Background | Genotype | Reference |
| --- | --- | --- | --- | --- |
| YKK200 | WT | BY4741 | *MATα his3Δ1 leu2Δ0 met15Δ0 ura3Δ0* | GE Dharmacon |
| YKK491 | Gal::Rps26 | BY4741 | *MATα NatMX6::pGAL1-Rps26A Rps26B::KanMX6 his3Δ1 leu2Δ0 met15Δ0 ura3Δ0* | (Ferretti et al 2017) |
| YKK1650 | ΔGID4 | BY4741 | *MATα gid4:: GID4::KanMX6 his3Δ1 leu2Δ0 met15Δ0 ura3Δ0* | GE Dharmacon |
| YKK1651 | ΔGID10 | BY4741 | *MATα gid4:: GID10::KanMX6 his3Δ1 leu2Δ0 met15Δ0 ura3Δ0* | GE Dharmacon |

**Table S2:** Plasmids used in this work.

| Plasmid | Description | Backbone | Reference |
| --- | --- | --- | --- |
| pKK3558 | TEF::Rps26A | pRS416 | (Ferretti et al 2017) |
| pKK31160 | TEF::Rps26A_P2S | pRS416 | This work |
| pkk30528 | Gal::Rps26A-HA | pRS426 | (Yang & Karbstein 2022) |
| pKK31150 | Gal::Rps26A_P2S-HA | pRS426 | This work |
| pKK31155 | Gal::Rps26A_D33N-HA | pRS426 | This work |
| pKK31157 | Gal::Rps26A_P2SD33N-HA | pRS426 | This work |
| pKK31177 | Gal::Rps26A_K28R-HA | pRS426 | This work |
| pKK31178 | Gal::Rps26A_K66R-HA | pRS426 | This work |
| pKK31179 | Gal::Rps26A_K70R-HA | pRS426 | This work |
| pKK31180 | Gal::Rps26A_K108R-HA | pRS426 | This work |
| pKK31181 | Gal::Rps26A_K116R-HA | pRS426 | This work |
| pKK31042 | Gal::Rps3-HA | pRS425 | This work |
| pKK31195 | TEF::Rps26A_K66R | pRS416 | This work |
| pKK31196 | TEF::Rps26A_K70R | pRS416 | This work |

